# Repetition Probability Effects for Chinese Characters and German Words in the Visual Word Form Area

**DOI:** 10.1101/2021.11.01.466762

**Authors:** Chenglin Li, Gyula Kovács

## Abstract

The magnitude of repetition suppression (RS), measured by fMRI, is modulated by the probability of repetitions (P(rep)) for various sensory stimulus categories. It has been suggested that for visually presented simple letters this P(rep) effect depends on the prior practices of the participants with the stimuli. Here we tested further if previous experiences affect the neural mechanisms of RS, leading to the modulatory effects of stimulus P(rep), for more complex lexical stimuli as well. We measured the BOLD signal in the Visual Word Form Area (VWFA) of native Chinese and German participants and estimated the P(rep) effects for Chinese characters and German words. The results showed a significant P(rep) effect for stimuli of the mother tongue in both participant groups. Interestingly, Chinese participants, learning German as a second language, also showed a significant P(rep) modulation of RS for German words while the German participants who had no prior experiences with the Chinese characters showed no such effects. Our findings suggest that P(rep) effects on RS are manifest for visual word processing as well, but only for words of a language with which participants are highly familiar. These results support further the idea that predictive processes, estimated by P(rep) modulations of RS, require prior experiences.

## 1. Introduction

Over the last two decades, the theory of predictive coding became a popular framework to understand how the brain encodes sensory information (Friston, 2005; Rao & Ballard, 1999). To achieve effective and precise encoding, the predictive brain constantly matches the incoming sensory information to prior expectations, thereby minimizing their mismatch (the predictive error) (Kok & de Lange, 2015).

Recently, predictive error has been associated with the well-known reduction of the neural responses for repeated presentations of a given stimulus (Desimone, 1996). Repetition suppression (RS) is a widely studied neural phenomenon when the repetition of a stimulus leads to the attenuation of the neuronal responses compared to its first presentation (Henson & Rugg, 2003). Despite the permanent interest in the topic the precise neural mechanisms of the effect are still unclear (for a review see Grill-Spector et al., 2006). In recent years, RS has been related to predictive coding as repeating a stimulus may induce its enhanced expectation and the reduction of prediction error, expressed in RS (for a review, see Grotheer & Kovács, 2016). Summerfield et al (2008) provided the first empirical evidence of the relationship between RS and predictive coding by modulating the statistical probability of stimulus repetitions [P(rep)] and using functional magnetic resonance imaging (fMRI). They compared the blood oxygenation level-dependent (BOLD) responses to face stimulus repetitions and alternations in two different contexts. A stronger RS effect was obtained in blocks, where the probability of repetition trials was higher when compared to blocks with less frequent repetitions. Later, the modulation of RS by P(rep) was confirmed in several subsequent studies using faces (Kovács et al., 2012; Larsson & Smith 2012). Interestingly, the P(rep) modulation of RS for non-face stimuli led to less consistent results. While some studies were able to find evidence of P(rep) modulations of RS for every-day objects (Mayrhauser et al., 2014), others failed to find it (Kovács et al., 2013). One possible explanation for these opposite results is that humans have different prior experiences with various stimulus categories, and this modulates the neural mechanisms, thereby the P(rep) effects of RS.

Indeed, it has been suggested that there is a tight connection between predictions and stimulus expertise, as gaining prior experiences leads to stronger associations among perceptual features or objects (for a review see Cheung and Bar, 2012). To explicitly test this hypothesis, Grotheer and Kovács (2014) adopted the paradigm of Summerfield et al., (2008), using two non-face stimulus categories with different levels of prior expertise: individual Roman letters and novel pseudo-letters. The results revealed P(rep) modulations on RS in the Letter Form Area (LFA; Thesen et al., 2012) for the well-known Roman letters, but not for the unfamiliar pseudo-letters, suggesting that prior experiences might indeed alter the underlying neural mechanisms of RS. However, it remains unclear whether this P(rep) effect (a) is observed for other stimuli, such as words as well; (b) can also be observed in other cortical areas than the LFA and (c) if learning a new stimulus category changes the P(rep) effects after a given training period or if life-long experiences are required, such as the one we all have with conspecific faces or literate persons have with characters of the familiar alphabet.

To address these questions, we performed an fMRI experiment, estimating the P(rep) modulation of RS for Chinese characters and German words in two participant populations: literate German participants having no experience with the Chinese language and native Chinese participants learning German as a second language for at least two years. We focused on the visual word form area (VWFA), located in the left occipitotemporal sulcus as previous studies suggested that this area reflects the visual and pre-lexical processes of word recognition (Cohen et al., 2000; Dehaene & Cohen, 2011). Our findings suggest that the P(rep) modulation of RS is manifest for visual word processing as well, but only if the language is known to the participants. These results support further the theory that the neural mechanisms of RS are modulated by prior experiences.

## 2. Results

### 2.1 Behavioral performance

German participants detected the target stimuli on average with 93.1 and 94.1% (±SE: 2.0 and 1.0 %) accuracy (average reaction time: 1143 and 1153 ms (±SE: 24 and 25 ms)) during the German word and Chinese character runs, respectively. The performance was not significantly different for the different blocks for either language. Chinese participants showed a similar pattern (average performance: 90.4 and 93.3% (±SE: 3.3 and 1.7 %); average reaction time:1117 and 1153 ms (±SE: 44 and 28 ms) during the German word and Chinese character runs, respectively) without any significant effect of blocks. In addition, no participant was aware of the distinct manipulations of P(rep) across the different blocks, based on their self-reported results after the fMRI measurement.

### 2.2 Neuroimaging results: German participants

#### 2.2.1 VWFA

A three-way repeated measures ANOVA revealed no significantly different activations for the German words and Chinese characters in the VWFA of the German participants (main effect of stimulus language: *F*(1, 19) = 0.02, *p* = 0.891, *η*_*p*_^*2*^ = 0.001).

We found a significant main effect of trial type for the words of the mother tongue of the German participants (2-way ANOVA with block and trial as factors; main effect of trials: *F*(1, 19) = 9.94, *p* = 0.005, *η*_*p*_^*2*^ = 0.34), with stronger BOLD signals in At (average BOLD signal ± SE: 0.169 ± 0.021) as compared to Rt trials (0.107 ± 0.028) (Figure 1). More importantly, the interaction of block and trial type was also significant for German words (*F*(1, 19) = 5.62, *p* = 0.028, *η*_*p*_^*2*^ = 0.23): post-hoc tests revealed a significant RS effect in the repetition (*Fisher post hoc test: p* = 0.005), but not in the alternation blocks (*p* = 0.869). This suggests that P(rep) modulates the magnitude of RS strongly for words of the mother tongue in the VWFA. This finding extends the previous findings of Grotheer & Kovács, (2014) who found similar strong interactions for Roman letters in the LFA of German participants.

**Figure 1.**
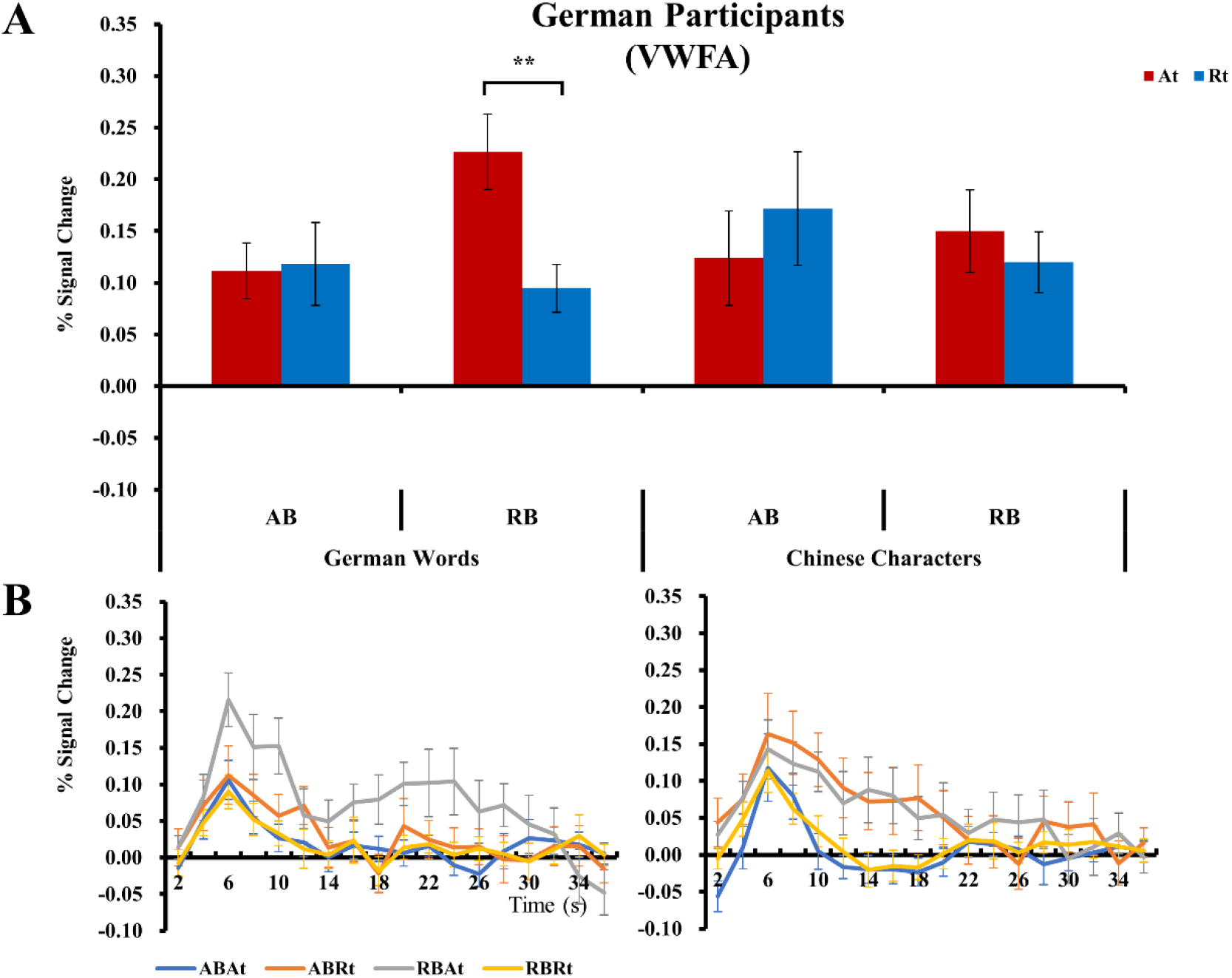
The BOLD results of the VWFA for German participants. ***A***, Average peak activation profiles of the VWFA for At and Rt trials for the German Words and for the Chinese Characters separately. ***B***, The time course of fMRI activity in the VWFA for German Words and for Chinese Characters. The hemodynamic response functions were derived from an FIR model with 2 s time bins. Error bars represent standard errors of means. **p < .01 (Fisher’s *post hoc* comparisons).

Interestingly, the same analysis showed no significant effects for Chinese characters (all *p values* ≥ 0.22), suggesting that neither repetition suppression, nor its modulation by P(rep) is present in the VWFA for the written symbols of an unknown language.

#### 2.2.2 LO

The BOLD signal in the LO was significantly stronger for Chinese characters, as compared to German words in the German participants (main effect of stimulus language: *F*(1, 19) = 43.45, *p* < 0.001, *η*_*p*_^*2*^ = 0.70). This is in line with those previous findings which found stronger activations in the LO for Chinese characters, as compared to French words in French participants (Szwed et al., 2014).

Area LO did not show either a significant main effect of hemisphere (*F*(1, 19) = 0.36, *p* = 0.554, *η*_*p*_^*2*^ = 0.02), or any interaction of hemisphere with stimulus, trial or block (all p values ≥ 0.166). For the German words the main effect of trial showed a non-significant tendency (*F*(1, 19) = 2.71, *p* = 0.116, *η*_*p*_^*2*^ = 0.13) and there was a significant main effect of block (*F*(1, 19) = 8.42, *p* = 0.009, *η*_*p*_^*2*^ = 0.31) (Figure 2). Similar to the VWFA, the block x trial interaction was also strongly significant in the LO (*F*(1, 19) = 10.39, *p* = 0.004, *η*_*p*_^*2*^ = 0.35) and post-hoc tests revealed a significant RS effect in the repetition (*Fisher post hoc test: p* = 0.0001), but not in the alternation blocks (*p* = 0.819). This suggests somewhat noisier, but similar processing of words of the mother tongue in the LO as in the VWFA.

**Figure 2.**
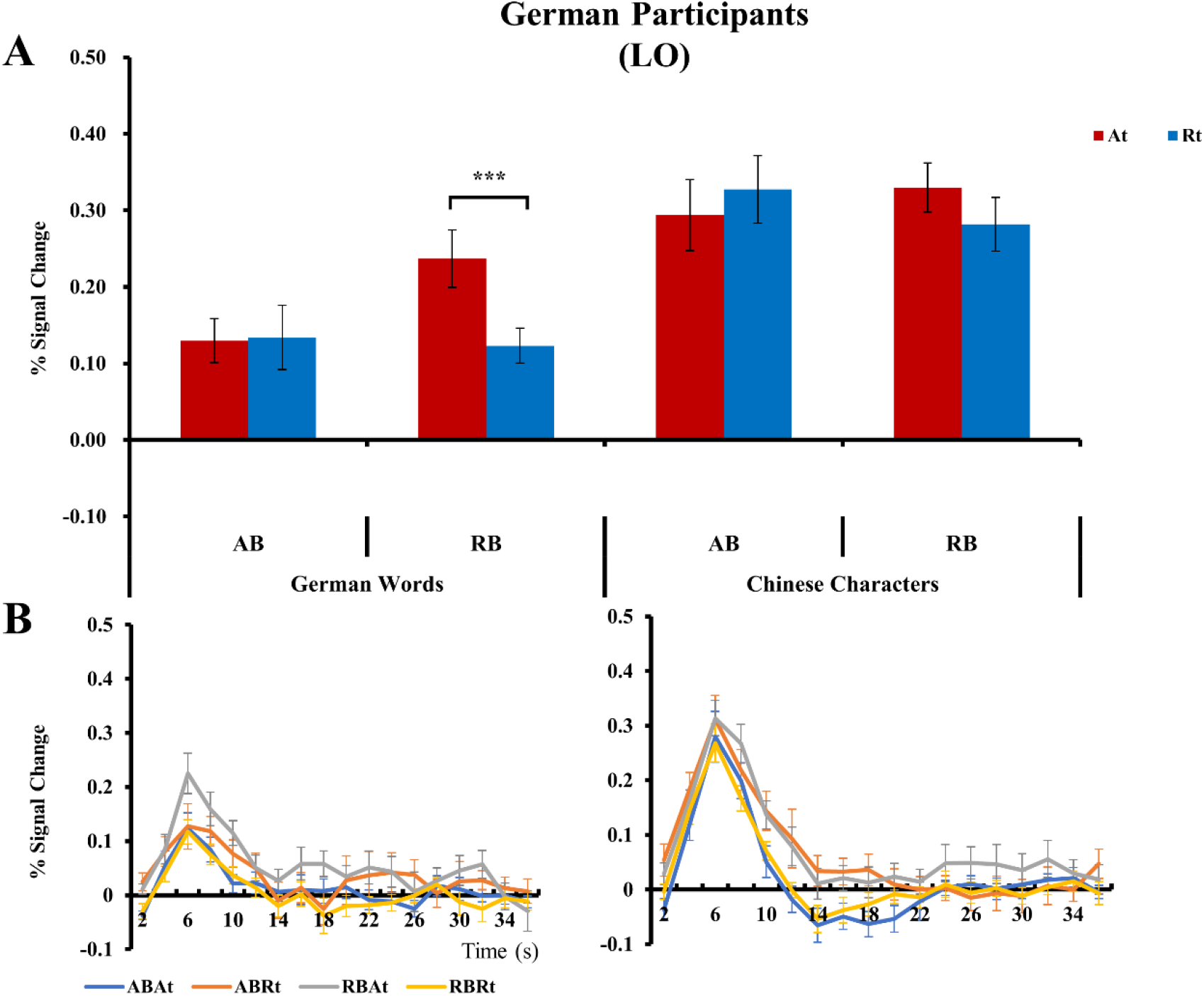
The BOLD results of the LO for German participants. ***A***, Average peak activation profiles of the LO for At and Rt trials for the German Words and for the Chinese Characters separately. ***B***, The time course of fMRI activity in the LO for German Words and for Chinese Characters. The hemodynamic response functions were derived from an FIR model with 2 s time bins. Error bars represent standard errors of means. ***p < .001 (Fisher’s *post hoc* comparisons).

For the Chinese characters neither the main effects of trial or block, nor their interaction approached significance (all *p value*s ≥ 0.21). This suggests that the LO, just like the VWFA, processes words of the mother tongue and an unknown language differentially.

### 2.3 Neuroimaging results: Chinese participants

#### 2.3.1 VWFA

The three-way repeated measure ANOVA revealed a significant main effect of stimulus language (*F*(1, 19) = 10.63, *p* = 0.004. *η*_*p*_^*2*^ = 0.36), with stronger BOLD signal for the Chinese characters (0.316 ± 0.046) as compared to the German words (0.191 ± 0.030) (Figure 3) in the Chinese participants.

**Figure 3.**
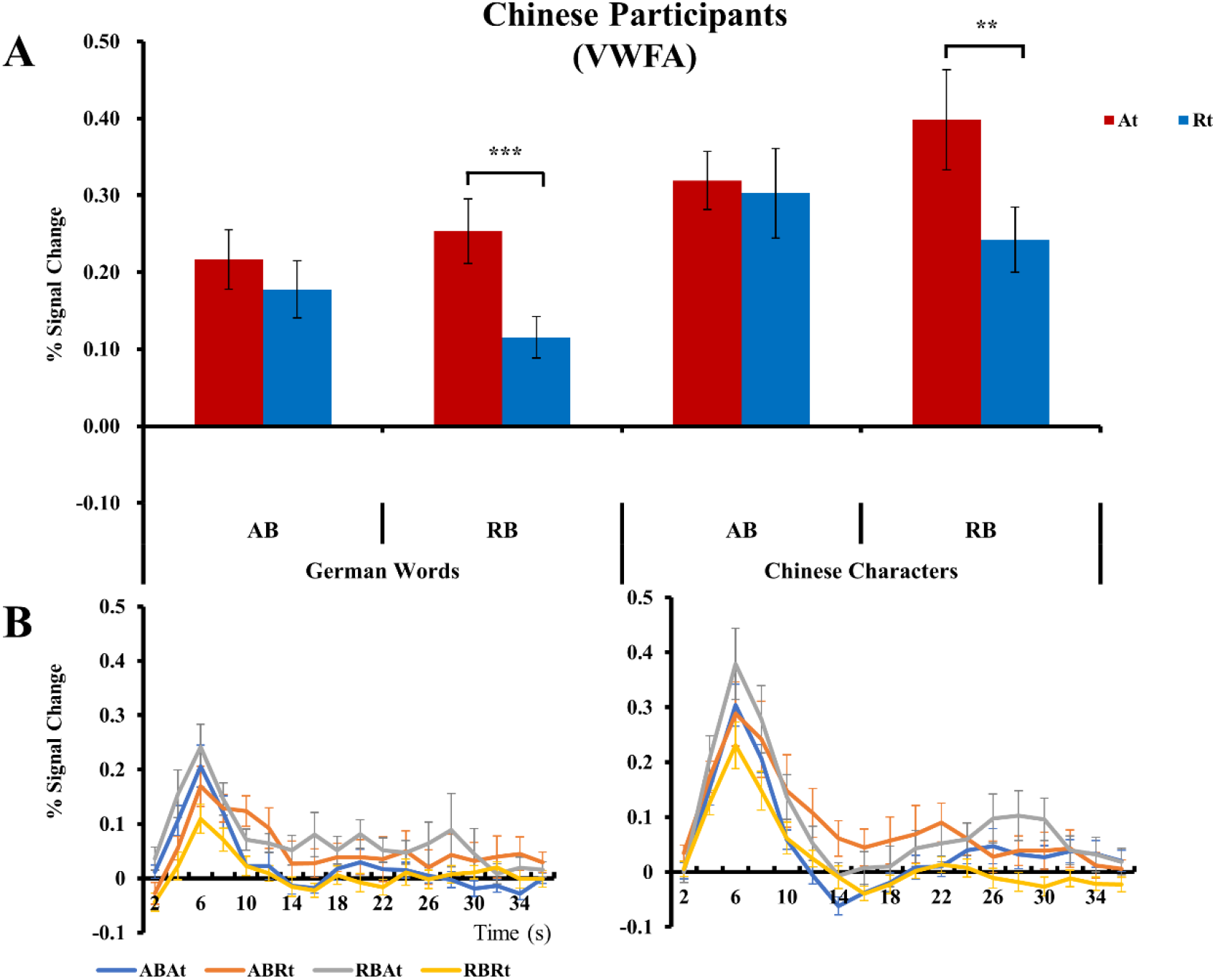
The BOLD results of the VWFA for Chinese participants. ***A***, Average peak activation profiles of the VWFA for At and Rt trials in the German Word and Chinese Character runs, respectively. ***B***, The time course of fMRI activity in the VWFA for German Word and Chinese Character runs, respectively. The HRF curves were derived from an FIR model with 2 s time bins. Error bars represent standard errors of means. **p < .01, ***p < .001 (Fisher’s *post hoc* comparisons).

Unlike for the German participants, in the Chinese we found significant RS effects in the VWFA both for German words (main effect of trial type: *F*(1, 19) = 18.51, *p* < 0.001, *η*_*p*_^*2*^ = 0.49) and for Chinese characters (main effect of trial type: *F*(1, 19) = 22.90, *p* < 0.001, *η*_*p*_^*2*^ = 0.55). More importantly, P(rep) modulations on RS were found for the characters of the mother tongue (block x trial for Chinese characters: *F*(1, 19) = 4.69, *p* = 0.043, *η*_*p*_^*2*^ = 0.20) as well as for the German words (block x trial: *F*(1, 19) = 6.72, *p* = 0.018, *η*_*p*_^*2*^ = 0.26). Post-hoc tests revealed that strong RS effects were observed in repetition blocks for both language stimuli (*Fisher post hoc test:* for German words: *p* = 0.000064, for Chinese characters: *p* = 0.0028), but not in the alternation block (*Fisher post hoc test:* for German words: *p* = 0.168, for Chinese characters: *p* = 0.719).

These results extend further the previous results with familiar Roman letters (Grotheer & Kovacs, 2014) and words (present study) in the sense that the characters/words of the mother tongue elicit both a significant RS in the relevant processing areas as well as its modulation by P(rep), independently of language and character-set. Importantly, the modulatory effects extend towards a second language as well, acquired by the participants only recently.

#### 2.3.2 LO

The activity of LO was significantly stronger for Chinese characters, as compared to German words in the Chinese participants as well (main effect of stimulus language: *F*(1, 19) = 55.52, *p* < 0.001, *η*_*p*_^*2*^ = 0.75). Area LO showed no main effect of hemisphere (*F*(1, 19) = 0.95, *p* = 0.341, *η*_*p*_^*2*^ = 0.05), or any interaction of hemisphere with stimulus, trial or block (all p values larger than 0.080) in the Chinese participants either. For the Chinese characters LO, just like the VWFA, showed a significant RS (main effect of trial: *F*(1, 19) = 5.66, *p* = 0.028, *η*_*p*_^*2*^ = 0.23) (Figure 4). Also mirroring the findings of the VWFA, the interaction of RS with P(rep) was also significant for the characters of the mother tongue of the Chinese participants (block x trial: *F*(1, 19) = 5.04, *p* = 0.037, *η*_*p*_^*2*^ = 0.21) and post-hoc tests revealed a significant RS effect in the repetition (*Fisher post hoc test: p* = 0.0068), but not in the alternation blocks (*p* = 0.888). For the German words LO showed a similar pattern. We observed a strong RS (main effect of trial: *F*(1, 19) = 8.89, *p* = 0.008, *η*_*p*_^*2*^ = 0.32) as well as an interaction that just exceeded the 0.05 p value threshold (block x trial: *F*(1, 19) = 4.14, *p* = 0.056, *η*_*p*_^*2*^ = 0.18) and post-hoc tests revealed a significant RS effect in the repetition (*Fisher post hoc test: p* = 0.017), but not in the alternation blocks (*p* = 0.782). Overall, the results obtained for the LO of Chinese participants are in line with those of the German participants in the sense that the LO and the VWFA seem to reflect similar P(rep) modulation of RS.

**Figure 4.**
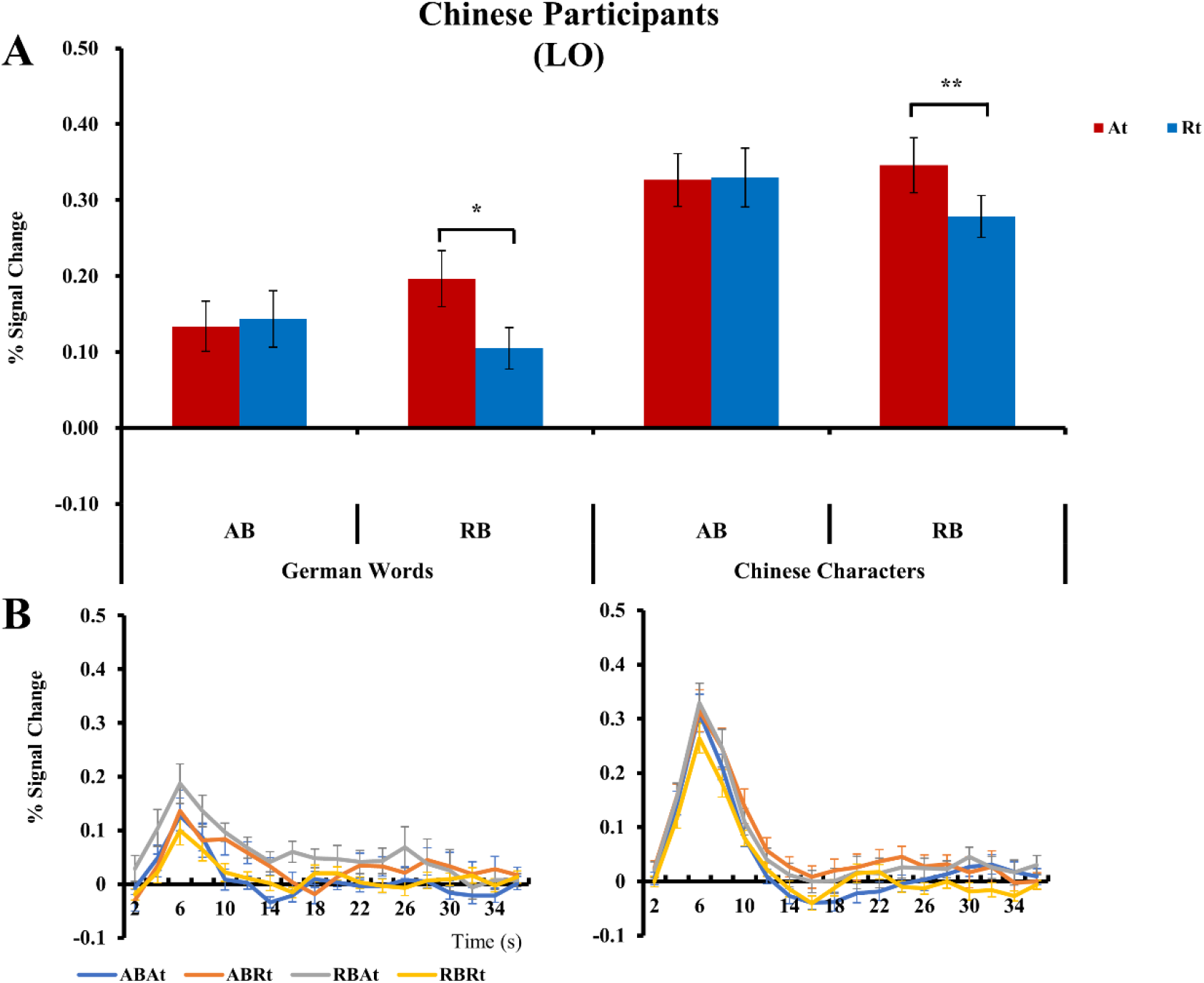
The BOLD results of the LO for Chinese participants. ***A***, Average peak activation profiles of the LO for At and Rt trials for the German Words and for the Chinese Characters separately. ***B***, The time course of fMRI activity in the LO for German Words and for Chinese Characters. The hemodynamic response functions were derived from an FIR model with 2 s time bins. Error bars represent standard errors of means. *p < .05, **p < .01 (Fisher’s *post hoc* comparisons).

### 2.4 Whole-brain analyses

To test further whether the effects of stimulus, repetition, or repetition probability can also be observed outside the predefined ROIs of our study we performed a group-level whole-brain analysis, separately for the German and Chinese participants. We applied a rigorous threshold of p < 0.05 (FWE corrected) with a minimum cluster size of > 50 voxels (Grotheer & Kovács, 2014). These analyses revealed a significant main effect of stimuli for both the German and Chinese participants: the activity of the middle occipital gyrus (*for average coordinates see Supplementary Material, Table 2*) was stronger for Chinese characters compared to German words. This is consistent with those previous studies which showed strong differences between the reading networks for Chinese and Latin alphabets (Szwed et al., 2014; Tan et al., 2005; Wu et al., 2012). No other statistically significant clusters were found when testing for the main effects of blocks, trials or their interaction. To make sure that no region is overlooked by the commonly applied, rigorous FWE corrected threshold we also used a more liberal (p < 0.001_uncorrected_) threshold to reanalyze our data. In this case the At > Rt contrast revealed one small cluster over the middle temporal gyrus (MNI coordinates: -58, -48, 8; cluster size: 10 voxels) in Chinese participants, suggesting repetition related response reduction. All statistically significant clusters of stimulus effect are listed in Supplementary Table 2.

## 3. Discussion

The main findings of the current study are the followings. (1) The VWFA and the LO show RS and its modulation by P(rep) for visually presented words/characters of the mother tongue but not an unknown language, in two different language populations. This result extends those previous findings which showed similar P(rep) effects within the fusiform face area and the letter form area. (2) The significant P(rep) effect for German words in Chinese participants and its lack for Chinese characters in Germans supports the idea that acquiring experience with a given stimulus category may be an important factor in determining P(rep) effects on RS. Further, it also suggests that learning a second language leads to changes of the neural mechanisms underlying RS in the VWFA.

To the best of our knowledge, the present study is the first to provide direct empirical evidence for the modulation of RS to written words by top-down probabilistic mechanisms. This result adds to the stimulus categories (Faces, Summerfield et al., 2008; Objects: Mayrhauser et al., 2014; Familiar Roman letters: Grotheer & Kovács, 2014) for which the significant P(rep) modulation of RS suggests the involvement of predictive coding mechanisms at least in human neuroimaging studies. Note however, that for every-day objects neither some neuroimaging nor non-human primate single-cell recordings found such effects (Kaliukhovich & Vogels, 2011; Kovács et al., 2013). Previously Grotheer and Kovács (2014) observed P(rep) effect in the LFA and LO for familiar Roman letters, but not for novel pseudo-letters. They suggested that an important factor in determining the emergence of P(rep) effect is the prior experience which the observer had with the stimuli. Indeed, the stimulus categories for which P(rep) modulation of RS was successfully shown are those where participants have typically extensive prior experience with (faces, words, and Roman letters). This conclusion is supported by the fact that prior experiences generally play a key role in forming predictions (Kok and de Lange, 2015).

The current study mainly focused on the VWFA, located in the left occipitotemporal sulcus and involved in the visual and prelexical processing of words (Cohen et al., 2000; Dehaene & Cohen, 2011). Previous studies have already shown significant RS in the VWFA both by neuroimaging (e.g., Barton et al., 2010; Glezer et al., 2015) and electrophysiological methods (e.g., Cao et al., 2015; Li et al., 2019; Mercure et al., 2011). Literate humans are considered as experts in the recognition of words of their mother tongue. Accordingly, the processing of written words of the mother tongue shows similarities to that of another stimulus categories of which humans are experts, for example familiar human faces (Davies-Thompson et al., 2016; for a review, see Moret-Tatay et al., 2020; Young and Burton, 2017). Therefore, it is not surprising that the VWFA is located over the occipito-temporal cortex (Cohen & Dehaene, 2004) in the close neighborhood of the fusiform face area (FFA; Kanwisher et al., 1997). In fact, it has been proposed that the VWFA emerges as the result of “cultural recycling” due to the potential re-use of neural structures previously related to face and object processing (Dehaene & Cohen, 2007). As a result, it has also been proposed that many structural, connectivity and functional properties of the VWFA are inherited from evolutionarily older brain circuits, such as the face-processing network (Hannagan et al., 2021). Thus, it is not surprising that similar P(rep) modulation of RS has been found within the VWFA for written words and in the FFA for faces (Summerfield et al., 2008).

Previous studies, using the identical paradigm as the present work have found P(rep) modulations on RS for familiar Roman letters, but not for novel pseudo-letters (Grotheer & Kovács, 2014). Therefore, it has been suggested that an extensive prior experience is necessary to the P(rep) effect to emerge. Here we compared the P(rep) effects on RS for the written words of two very different languages as mother tongue. Thus, our results not only extended the P(rep) effects to visual word processing, but also show the same P(rep) effects in two separate participant groups and for two very different lexical stimuli: words composed of letters of the German alphabet and Chinese characters, which are logograms, representing words or morphemes. Our results are also in line with previous studies which found similar RS effects both for the written words of alphabetic and logographic languages (Cao et al., 2014; Maurer et al., 2005). Importantly, the current results also suggest that the P(rep) modulation of RS is a universal effect, which develops across languages and cultures and is independent of the alphabetical or logographic nature of the written characters.

Another result of the current study is that the Chinese participants showed a significant P(rep) effect on RS for German words, which they study as a second language for several years. Explicit learning is an obvious way to obtain experience and knowledge with something. For instance, perceptual learning can improve various perceptual abilities, ranging from discriminating visual features (for example contrast, orientation or shape) (Schoups et al., 1995; Yu et al., 2004; for a review, see Bi & Fang., 2013), to object recognition capacities (Op de Beeck & Baker, 2010). Previous studies indicated that the human visual system is generally more sensitive to learned compared to novel objects (Grill-Spector et al., 2000; Sigman et sl., 2005; Op de Beeck et al., 2006). For instance, Song et al. (2010) trained two groups of participants, either in a visual association or a shape discrimination task. Their results showed a stronger learning effect in the VWFA as compared to the LO when trained in the association learning tasks, but a greater learning effect in the LO when trained in the shape discrimination task. This suggests that the VWFA is sensitive to top-down learning contexts during the encoding of objects. Our results suggest that the top-down effects, elicited by repetition probability modulations of stimulus blocks, are formed only after long-term accumulation of experiences with the stimuli.

One limitation of the current study is its unbalanced nature: we tested native German speaker, who were entirely naïve regarding Chinese characters as well as native Chinese speakers who were also exposed to German words in the last years. Ideally, in a fully balanced design, we should have tested Chinese participants who were never exposed to German words and Latin characters as well as Chinese-speaking German participants. On the one hand, as the study was conducted in Germany it was impossible to find Chinese participants who are entirely naïve regarding German words. In fact, in the light of the current educational system in China, where children are almost always exposed to western languages (and thereby to Latin characters) during their education we feel that the testing of such a population is not feasible in the future either. On the other hand, due to the scarcity of Chinese-speaking German participants in our experimental participant population (we only could record 5 native German participants, who learned Chinese for at least 2 years until the end of the current experiment) their data was also not analyzed here. However, two arguments suggest that the current data suggests the role of experience in modulating the neural mechanisms of RS. First, we found P(rep) modulations on RS for two fundamentally different languages, one with alphabetic and another with logographic orthographies, suggesting its independence of the writing system. Second, the modulation effect transferred across writing systems, as the findings for German words in Chinese participants show. Nonetheless, for a full and language-independent support of the conclusion that P(rep) modulations of RS require experiences with the words of a particular language one should replicate the current experiment with a fully balanced design ideally in two different native populations with and without experiences with the other language. Alternatively, a longitudinal training study could also be performed in the future to investigate the modulation of the P(rep) effect during the learning of a foreign language.

In addition to the VWFA, the LO (Malach et al., 1995) is commonly considered as a key area involved in object processing, but is also responsive to faces (Lerner et al., 2001). The present study shows that the LO responds to German words and Chinese Characters as well. Previous studies found a significant modulation of P(rep) on RS in the area for various stimulus categories (faces: Kovács et al., 2012; Larsson & Smith, 2012; Familiar Roman letters: Grotheer & Kovács, 2014), but not for novel pseudo-letters. Consistent with these findings, the present study shows that the characters/words of the mother tongue induce a significant P(rep) modulation on RS in the LO, and these modulatory effects extend towards a second language acquired by the participants relatively recently. This indicates that P(rep) effects on RS in LO are not specific to one stimulus category, and that they depend on the prior experiences with the stimuli, such as the letters and words of the familiar alphabet and language for literate persons. Interestingly, some previous studies using the hemifield change paradigm found that the VWFA and LO reflect different sensitivity for repetition suppression in word recognition (Strother et al., 2016, 2017; Zhou et al., 2019). For instance, Zhou et al (2019) found that the VWFA show both font-sensitive and font-invariant effects, while the LO reflects greater font sensitivity. As the modulatory pattern of prior experiences in the LO was similar to that in the VWFA, our current results suggest that the LO and the VWFA reflect similar neural processing stages.

We argue that that written word processing is an optimal cognitive process to test the validity of the idea of the predictive brain from an experimental perspective. Generally, word recognition includes visual, auditory as well as semantic processes. Previous studies suggested that predictive mechanisms are important in determining the RS, observed in the LFA for simple graphemes or letters (Grotheer & Kovács, 2014), in the superior temporal gyrus for word stress processing (Honbolygó et al., 2020) and in the superior temporal gyrus for lexical representations (Wang et al., 2021) as well as for semantic processes (Qin et al., 2021). These results combined with the current study, in which the involvement of predictive processes is suggested for visual word recognition in the VWFA, suggest that top-down probabilistic prediction effects are reflected across several stages of language processing. Future studies should address this issue in a more systematic manner by using for example artificial language learning.

In conclusion, the present study shows that top-down effects, expressed in probabilistic manipulations of RS can be observed in the VWFA/LO and this effect is only manifest for words of a language with which participants have extensive prior experiences. These findings support the idea that predictive processes, measured by P(rep) modulation, require extensive prior experiences.

## 4. Methods

### 4.1 Participants

Twenty healthy German participants with normal or corrected-to-normal vision took part in the experiment (15 females; mean age = 22.4 years; SD = 3.6 years; four left-handed). These participants were native German speakers and had no experience with Chinese language.

Twenty-two healthy Chinese participants were also recruited who were living in Germany. Two of these were excluded from the analysis due to their poor German knowledge as they recognized only 20% and 30% of the test words. The remaining 20 participants (12 females; mean age = 26.2 years; SD = 2.7 years; one left-handed) had normal or corrected-to-normal vision. On average, they have studied German for more than 4 years (M ± SD = 4.8 ± 2.1 years).

The sample size was determined, based on previous studies testing the same P(rep) effects for other stimulus categories and finding significant effects for faces (Grotheer & Kovács, 2014; Summerfield et al., 2008). Participants received partial course credits, monetary compensation, or their own 3D-printed brain-model as a compensation. All participants were informed about the experimental procedures and completed a written consent prior to the experiment, and they had no history of any neurological disorders. The research protocol was approved by the ethics committee of the Friedrich-Schiller-Universität Jena and conducted in accordance with the guidelines of the Declaration of Helsinki.

### 4.2 Stimuli and task

The experimental design of the present study was identical to the previous studies of our laboratory (Grotheer & Kovács, 2014), with the exception that 240 high-frequency German words and 240 high-frequency Chinese Characters were used as stimuli. The German words were selected from an online German linguistic database (dlexDB (available at www.dlexdb.de); Heister et al., 2011), ranging in length from 4 to 7 letters and were presented with a height of 1.2° visual angle from a viewing distance of 100 cm. The Chinese characters were selected from the modern Chinese frequency dictionary (Wang et al., 1986) with the number of strokes varying between 4 and 8 and subtending angles of 4.8° × 4.8° from a viewing distance of 100 cm. The German words and Chinese characters were presented in standard typefaces (Sans Serif and Simhei fonts, respectively; for examples see Figure 5A). No significant difference in word frequency between German words (*M* = 0.59/1000, range: 0.14/1000 – 4.93/1000) and Chinese characters (*M* = 0.61/1000, range: 0.17/1000 – 1.19/1000; *t* (478) = 0.408, *p* = 0.683) was present. All stimuli were presented centrally on a uniform gray background, back-projected via an LCD video projector onto a translucent circular screen. Psychtoolbox (Version 3.0.15) was used for stimulus presentation and behavioral response collection, controlled by Matlab R2013a (The MathWorks).

**Figure 5.**
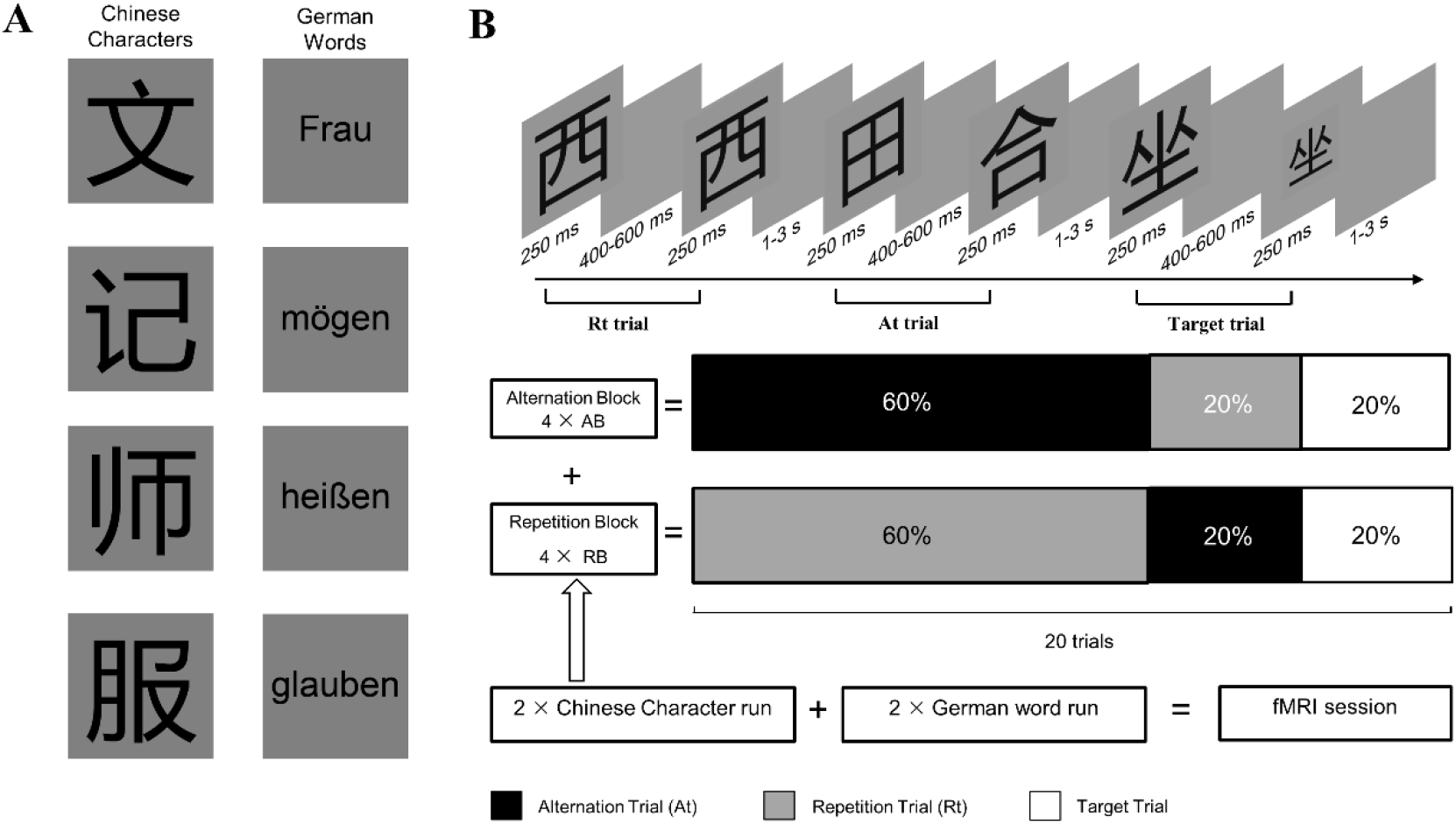
***A***, Examples of the words stimuli (Left column, Chinese Characters. Right column, German Words) used in the experiment. ***B***, Schematic illustration of the experimental design and task procedures. Rt-repeated trials, At-alternation trials, AB: alternation block, RB: repetition block.

The experimental procedures were identical to Grotheer & Kovács (2014), except for stimuli. Participants were asked to complete four experimental runs, two with German words and two with Chinese character stimuli, in a counterbalanced order. In each trial (Figure 5B), a stimulus-pair was presented (exposition time: 250 ms each), separated by an inter-stimulus-interval (ISI) of 400-600 ms (randomized across trials), and randomly followed by an Inter-trial-interval (ITI) of 1,000 ms or 3,000 ms. The first stimulus (S1) was either identical to (Repetition trials, Rt) or different from the second one (S2; Alternation trials, At). In addition, two different types of blocks were created, each repeated four times within a single run. In the Repetition Blocks (RB), 75% of the trials were Rt while 25% were At. In contrast, the Alternation Blocks (AB) were composed of 75% At and 25% Rt (Summerfield et al, 2008). Therefore, each run included 160 trials of the four different conditions (ABAt, ABRt, RBAt, and RBRt) in a randomized order. To avoid local feature adaptations, the size of either S1 or S2 (chosen randomly) was reduced by 18% during each trial. The design and parameters of the German word and Chinese character runs were identical, except for the stimuli. Participant’s task was to maintain central fixation and signal the occurrence of stimuli which were 60% smaller than the rest. Such target trials composed 20% of all trials (equally for Rt and At trials) and were modelled explicitly during the analysis but were not analyzed further.

### 4.3 Imaging parameters and data analysis

Imaging data were collected on a 3-Tesla MRI scanner (Siemens MAGNETOM Prisma, Erlangen, Germany), with a 20-channel head coil. T2*-weighted images were collected with the following parameters: FOV = 204 × 204 mm^2^; 34 slices; TR = 2000 ms; TE = 30 ms; flip angle = 90 deg; voxel size: 3 × 3 ×3 mm^3^. High-resolution T1-weighted images (192 slices; TR = 2300 ms; TE = 3.03 ms; flip angle = 90 deg; voxel size: 1 × 1 ×1 mm^3^) were acquired to obtain 3D structural scans with the same 20-channel head coil. Data were pre-processed using SPM12 (Welcome Department of Imaging Neuroscience, London, UK). Briefly, the functional images were slice-timed, realigned, co-registered to structural scans. The functional images were normalized to the MNI-152 space, resampled to 2 × 2 × 2 mm resolution, and spatially smoothed using an 8-mm Gaussian kernel.

Regions of interests (ROIs) were defined by additional functional localizer runs. Chinese characters, German words, line drawings of objects, and Fourier-randomized noise patterns were presented (230 ms exposure time; 20 ms inter-stimulus-interval) in blocks of 10 s, interrupted by breaks of 10 s and repeated four times. Each block included 40 images, which the size of 480x 480 pixels with a grey background. Participants were simple asked to pay attention to the stimuli and maintain fixation.

As previous studies, applying similar paradigms found that for faces the FFA while for Roman letters the LFA showed the largest effects (Grotheer & Kovács, 2014; Kovács et al., 2012, 2013; Summerfield et al., 2008), here we concentrated on the VWFA and the lateral occipital complex (LO; Malach et al, 1995). For the German participants the VWFA was localized individually by contrasting German words with noise images and its location was established as the local maximum from the t-maps with a threshold of *p*_FWE_ < 0.05 (n = 11) or p < 0.001 uncorrected (n = 9) on the single-subject level. The average MNI coordinates (±SE) for the VWFA were -44.8 (1.3), - 65.0 (2.6), -11.0 (1.1). The individual coordinates are listed in the Supplementary Table 1. In addition to the VWFA, we also estimated the BOLD signal in the LO as previous studies showed strong and significant p(rep) effects in this area for Roman letters (Grotheer & Kovács, 2014). The LO [average MNI coordinates (±SE): 42.1(1.3), - 78.0(1.5), -3(1.4) for the right, and -41.7(1.2), -78.9(1.4), -5.3(1.6) for the left hemisphere] was localized individually by contrasting line drawing of objects with noise images and the location was established as the local maximum from the t-maps with a threshold of *p*_FWE_ < 0.05 on the single-subject level.

For the Chinese participants the VWFA was localized by contrasting Chinese characters with noise images individually and the location was established as the local maximum from the t-maps with a threshold of *p*_FWE_ < 0.05 (n = 13) or p < 0.001 uncorrected (n = 7) on the single-subject level. The mean MNI coordinates (±SE) for the VWFA were -43.6 (0.9), -65.7 (2.6), -13.8 (1.0). The coordinates of the VWFA did not differ significantly across participant groups (X coordinate: *t*(38) = 0.448, *p* = 0.656; Y coordinate: *t*(38) = 0.205, *p* = 0.838; Z coordinate: *t*(38) = 2.015, *p* = 0.051). The LO [average MNI coordinates (±SE): 41.8(1.2), -79.0(1.3), -4.3(1.6) for right, and - 42.3(1.1), -78.7(1.0), -4.7(1.4) for left hemisphere] was localized individually by contrasting line drawing of objects with noise images and the location was established as the local maximum from the t-maps with a threshold of *p*_FWE_ < 0.05 on the single-subject level. The coordinates of the LO was also similar across participant groups (all *p* values ≥ 0.539). Canonical hemodynamic response functions were extracted using MarsBaR 0.44 toolbox for SPM 12 (Brett et al., 2002). The group-level ROIs in German and Chinese participants were visualized using xjView toolbox (http://www.alivelearn.net/xjview) and the average locations of VWFA and LO are shown in Figure 6, separately for German and Chinese participants. The analysis of these coordinates (between-subject t-tests, separately for x, y and z coordinates; *p* > 0.05 for every comparison) revealed that despite the fact that we used different contrasts to determine the location of these areas the VWFA and LO of both groups are overlapping in the two participant groups.

**Figure 6.**
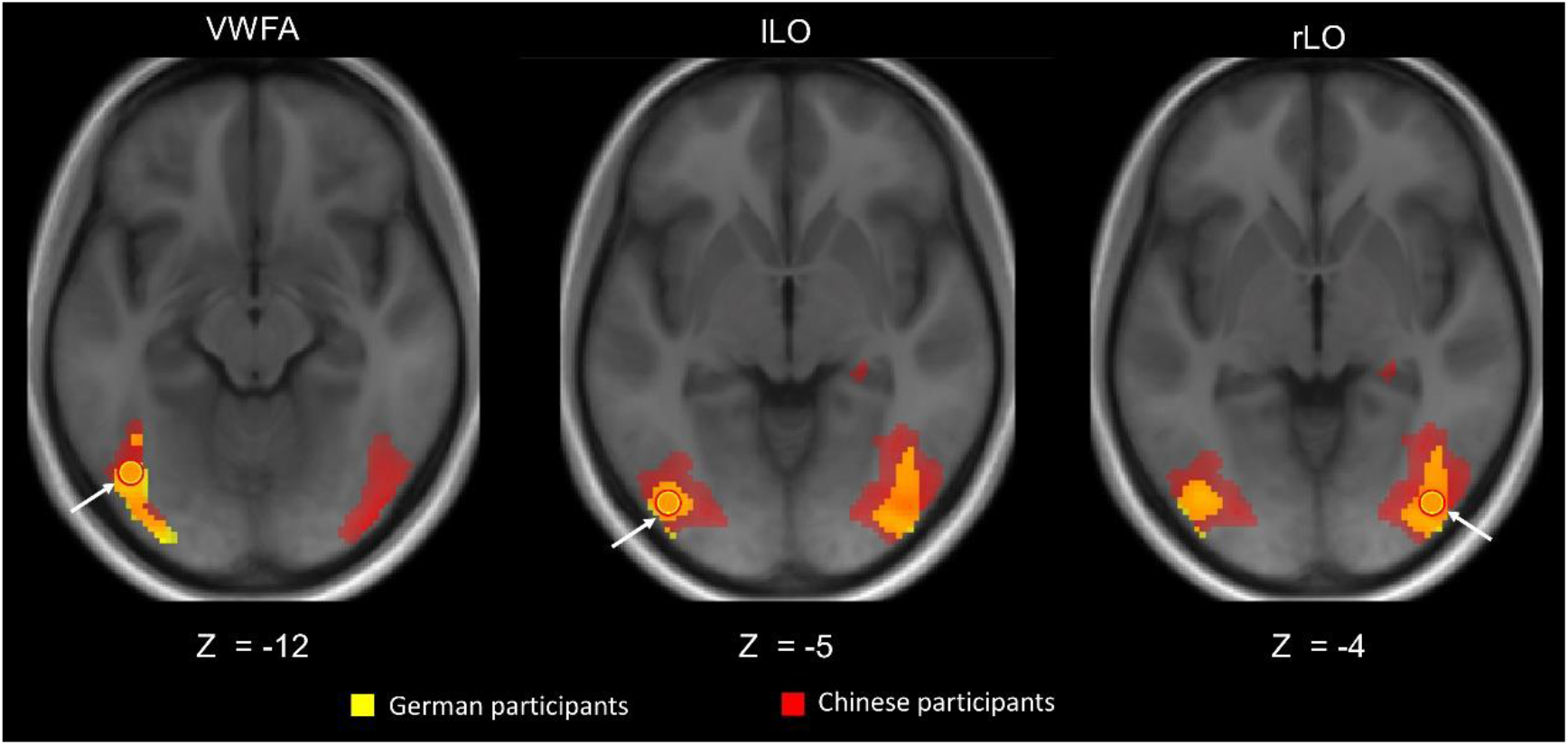
Group-level ROIs in German and Chinese participants. Yellow and red areas indicate the results in German and Chinese participants, respectively. The circles mark the average coordinates for both groups.

To directly compare with previous studies that used the same paradigm to investigate the repetition probability effects for objects and letters (Grotheer & Kovács, 2014; Kovács et al., 2013), the peak BOLD values were extracted from the event-related runs for the non-target trials in the VWFA and were analyzed by repeated-measures ANOVAs with stimulus language (German words vs. Chinese characters), block type (AB vs. RB) and trial type (At vs. Rt) as within-subject factors, separately for the German and Chinese participants. Additionally, since the general shape and visual complexity of the German words and Chinese characters are different (Tan et al., 2001), we performed another 2 × 2 repeated-measures ANOVAs with block type (AB vs. RB) and trial type (At vs. Rt), separately for German words and Chinese characters in the German and Chinese participants. For the LO, we conducted repeated-measures ANOVA with hemisphere (left vs. right), stimulus language (Chinese Characters vs. German words), block (AB vs. RB) and trial type (At vs. Rt) as within-subject factors, separately for the German and Chinese participants. All multiple comparisons of post-hoc tests were corrected by the Fisher’s method.

## Acknowledgements

This work was supported by a Deutsche Forschungsgemeinschaft Grant [KO3918/5-1]. Chenglin Li was supported by the China Scholarship Council (CSC) scholarship (201808330399) during the study. The authors thank Sophie-Marie Rostalski for her comments on the paper; Marie Wächter and Wenbo Wang for their help in the stimuli collection and participant recruitment.

