## Supplemental Tables 1 and 2 for "Repetition Probability Effects for Chinese Characters and German Words in the Visual Word Form Area"

Supplementary Table 1. Individual-participant VWFA and LO coordinates in German and Chinese participants

| Participants | Number | VWFA | Left LO | Right LO |
| --- | --- | --- | --- | --- |
| German | S01 | -42, -64, -10 | -44, -76, -10 | 42, -62, -16 |
|  | S02 | -40, -58, -4 | -38, -78, -4 | 40, -70, -10 |
|  | S03 | -48, -60, -18 | -40, -82, 4 | 46, -80, 2 |
|  | S04 | -38, -48, -14 | -44, -76, -2 | 42, -78, -2 |
|  | S05 | -46, -52, -12 | -36, -86, 10 | 38, -72, -8 |
|  | S06 | -38, -68, -2 | -50, -78, -4 | 44, -84, 4 |
|  | S07 | -56, -66, -10 | -34, -84, -18 | 38, -84, -2 |
|  | S08 | -54, -46, -10 | -42, -70, -8 | 54, -72, 0 |
|  | S09 | -44, -86, -6 | -46, -84, -6 | 40, -82, -2 |
|  | S10 | -42, -70, -10 | -48, -78, -2 | 38, -88, -4 |
|  | S11 | -44, -76, -14 | -54, -78, 4 | 52, -76, 2 |
|  | S12 | -56, -66, -10 | -36, -86, -14 | 36, -70, -12 |
|  | S13 | -42, -76, -10 | -38, -66, -12 | 46, -78, 4 |
|  | S14 | -42, -58, -20 | -44, -90, -2 | 40, -86, -2 |
|  | S15 | -44, -68, -12 | -36, -66, -10 | 40, -70, -8 |
|  | S16 | -38, -82, -4 | -40, -80, -6 | 38, -78, -4 |
|  | S17 | -42, -48, -18 | -36, -82, -14 | 30, -86, -2 |
|  | S18 | -50, -76, -10 | -42, -78, -8 | 44, -80, -12 |
|  | S19 | -44, -76, -14 | -38, -84, -6 | 48, -82, 8 |
|  | S20 | -40, -56, -12 | -48, -76, 2 | 46, -82, 4 |
| Chinese | S01 | -42, -54, -18 | -40, -82, -12 | 42, -76, -12 |
|  | S02 | -44, -50, -20 | -40, -80, -2 | 42, -78, -6 |
|  | S03 | -42, -52, -12 | -44, -76, 2 | 52, -82, -2 |
|  | S04 | -46, -56, -22 | -52, -72, 0 | 44, -74, -14 |
|  | S05 | -46, -68, -12 | -40, -84, 0 | 44, -84, -2 |
|  | S06 | -42, -64, -14 | -30, -88, 4 | 46, -76, -16 |
|  | S07 | -42, -74, -16 | -42, -84, -12 | 42, -82, -6 |
|  | S08 | -46, -68, -8 | -48, -78, -12 | 44, -78, -4 |
|  | S09 | -38, -83, -13 | -42, -80, -10 | 40, -84, 8 |
|  | S10 | -46, -58, -10 | -40, -80, -2 | 42, -78, -6 |
|  | S11 | -42, -78, -6 | -44, -76, -2 | 34, -76, 6 |
|  | S12 | -48, -78, -10 | -50, -76, -10 | 42, -66, -2 |
|  | S13 | -46, -54, -14 | -40, -74, 6 | 44, -76, 6 |
|  | S14 | -38, -78, -16 | -42, -72, -4 | 40, -74, -4 |
|  | S15 | -46, -76, -12 | -38, -74, -4 | 40, -86, -4 |
|  | S16 | -36, -82, -16 | -36, -80, -16 | 30, -92, 0 |
|  | S17 | -54, -72, -8 | -44, -80, -4 | 48, -76, -10 |
|  | S18 | -40, -50, -18 | -40, -80, -6 | 42, -82, 2 |
|  | S19 | -44, -54, -18 | -48, -76, 2 | 32, -76, -16 |
|  | S20 | -48, -66, -14 | -46, -82, -12 | 46, -84, -4 |

Supplementary Table 2. Summary of significant activations identified from the group level whole-brain analysis

| Contrast | Region | Hemisphere | German Participants | | Chinese Participants | | Threshold |
| --- | --- | --- | --- | --- | --- | --- | --- |
|  |  |  | MNI Coordinates | Cluster size (Voxels) | MNI Coordinates | Cluster size (Voxels) |  |
| Chinese Characters > German words | Middle occipital gyrus | R | 28, -88, 12 | 2519 | 26, -90, 8 | 1152 | P < 0.05, FWE |
|  | Middle occipital gyrus | L | -16, -98, 6 | 1533 | -18, -94, 6 | 555 | P < 0.05, FWE |
|  | Inferior occipital gyrus | L | - | - | -44, -78, -8 | 273 | P < 0.05, FWE |
| At > Rt | Middle temporal gyrus | L | - | - | -58, -48, 8 | 10 | P < 0.001 uncorrected |
